# Methodological comparison of different approaches to cultivate postnatal organotypic hippocampal cultures

**DOI:** 10.1101/2022.05.11.491471

**Authors:** Vivien Czapla, Pia Reiterer, Klaus Funke

## Abstract

To study molecular and cellular processes of the brain *in vitro*, the cultivation of postnatal organotypic hippocampal slices should be established. The methodological approach had to be optimized several times because the result of the cultivation was not optimal. Initially, cultivation was performed by roller tube culture, which is the first developed cultivation method for these cultures. As the cultures detached or even dissolved, further cultivation was carried out using inserts, which represent a modification of the roller tube cultures. In the further course it came also to the use of different preparation methods and to the exchange of the medium and the incubator, in order to obtain a result of the organ cultivation corresponding to the expectations. The results show that cultivation on semi-permeable membranes has an advantage over roller tube cultures. On the one hand, there is no detachment and dissolution of hippocampal slices due to the mechanical influence and on the other hand, the tissue shows an improvement in general condition compared to roller tube cultures. Furthermore, it was found that an acceleration of the preparation time has positive effects on the neurons. In addition, it could be shown by the recorded time series that a cultivation over a longer period of time stabilizes the tissue. If the hippocampi remain connected to the entorhinal cortex, wallerian degeneration does not occur, resulting in longer survival of the cells in the postnatal cultures and thus strengthening the tissue.

## 1 INTRODUCTION

Organ culture models serve as a promising alternative to animal models, as in vitro cultivation reduces the number of experimental animals. By preserving the tissue in its three-dimensional compsition, it is possible to analyse structural and synaptic organizations (Gstraunthaler und Lindl 2013; Humpel 2018). The viability of organotypic postnatal hippocampal cultures, as well as other organs, can be preserved over a limited period of time (Gogolla et al. 2006). Suitable conditions for organ cultures are important starting with preparation. By cooling down the tissue the life span of the organs can be extended after removal of the organism. Appropriate ambient temperature, proper air and gas mixture ratio, and regular medium replacement also play an important role in culturing (Gstraunthaler und Lindl 2013). Organotypic brain sections were first cultured in cell culture tubes by Gähweiler in 1981, whereas the methodology was described by Hogue in 1947 (Gähwiler 1981a, 1981b; HOGUE 1947). Brain sections are attached to coverslips using plasma and thrombin and placed in cell culture tubes. During cultivation the tubes rotate so that the tissue periodically comes into contact with either medium or air. The problem with this method is that the cells disorganize, die, or disintegrate over time (Humpel 2015). In 1991, cultivation on semi-permeable membranes was introduced by Stoppini et al. (Stoppini et al. 1991). Brain slices are placed on the membrane and receive their nutrients at the interface of membrane and medium. In this case, the tissue is not covered by the medium. An advantage of this cultivation is that no tissue sections can be lost and there is mechanical stress only during medium exchange (Humpel 2015).

## 2 EXPERIMENTAL PROCEDURES

### 2.1 Animals

The experimental animals were ordered from Charles River Laboratoriers (Sulzfeld, Germany). Pregnant female Wistar rats (Clause 1993) were delivered on day E13. The rats were kept in an open shelf system in type IV cages with a floor area of 800 cm2. The dams were kept individually. The room temperature was between 21 - 23 °C and the day-night rhythm was 12 hours. Lights were on from 6 am - 6 pm. Water and food were available to them *ad libitum*. The selection of juveniles was randomized and sex-independent. The age was between P5 and P10.

### 2.2 Medium preparation

Since two different preparation solutions and cultivation media were used, a distinction is made between the Hanks solution (Sigma-Aldrich) and the Minimum Essential Media (MEM) (Gibco 21575-022) solution and the Basal Medium Eagle (BME) (Gibco 41010-026) medium and the MEM medium. The Hanks solution was prepared consisting of 500 ml Hanks’ Balanced Salt Solution, 5% glucose (Sigma 68769) and 100 U/ml penicillin-streptomycin (Sigma P0781). Before usage it was bubbled with carbogen. The MEM solution was prepared consisting of 470.5 ml MEM, 5 ml Glucose 45 %, 12.5 ml HEPES 1 M (Invitrogen 15630-056), 5 ml Penicillin-Streptomycin and 2 ml NaOH 1 N. It was not bubbled with carbogen and the pH was adjusted between 7.3 and 7.4. The BME medium is composed of 25% heat-inactivated Horse Serum (Gibco 26050-088), 25% HBSS, 10% Glucose Stock Solution, 0.6 µg/ml NGF (Sigma), 200 mM L-Glutamine(Sigma), 100 U/ml Penicillin-Streptomycin and 0.2% Phenol Red. The mixture was then made up to 100 ml with BME. The MEM medium was composed of 42% MEM, 25% BME, 25% heat-inactivated Horse serum, 25 mM HEPES buffer (Invitrogen 15630-056), 0.2% sodium bicarbonate (Invitrogen 25080-060), 100 U/ml penicillin-streptomycin, and 2 mM glutamax (Gibco 35050-038). Both media were heated in a water bath before each use. The MEM medium was additionally adjusted to a pH of 7.3 after heating.

### 2.3 Preparation of the Hippocampus

All utensils and materials used were refrigerated. During dissection, the animal (between P8 and P10) was first decapitated with scissors and the head was transferred to a Petri dish containing frozen Hanks solution (Vibratome (first approach) and Chopper). The subsequent dissection procedure differed, depending on usage of either a vibratome or a chopper.

#### Vibratome (VT1000S, Leica Mikrosysteme Vertrieb GmbH, Wetzlar) (first approach)

The brain was carefully exposed on ice with small scissors and tweezers and transferred to 4 °C cold Hanks solution enriched with carbogen. After one minute, the brain was again spooned onto a small Petri dish containing frozen solution, which was lined with filter paper and a little Hanks solution. A scalpel was then used to cut off as much brain tissue as possible around the hippocampus. The hemispheres were separated by a cut and both attached to a cooled plate using tissue glue which was then attached to the vibratome in a buffer dish. The buffer dish was filled with ice-cold Hanks solution and cooled with ice from the outside. 350 µm thick brain sections were made along the sagittal plane. These brain sections were soaked up with a disposable pipette and transferred to a separate Petri dish containing cooled Hanks solution. Hippocampi were then carefully dissected out under a microscope by removing the surrounding tissue. The hippocampi were then transferred to a new Petri dish containing fresh, chilled Hanks solution.

#### Chopper (McIlwain Tissue Chopper, Plano GmbH, Wetzlar)

The brain was carefully exposed with small scissors and forceps, as much brain tissue as possible was separated around the hippocampus with a scalpel, and the hemispheres were separated. Hemispheres were then transferred to a Petri dish containing 4 °C cold carbogenan-enriched Hanks solution. In the next step, the striatum, thalamus, midbrain, and brainstem were carefully removed with forceps, exposing the cortex with the hippocampus attached. The hippocampus was then rolled out and separated from the cortex. The hippocampi of both hemispheres were placed longitudinally on the cut surface of the chopper and the fluid adjacent to the tissue was removed with a filter paper. The thickness of the hippocampal sections was 350 µm. After cutting, the tissue was carefully transferred with a spatula to a new Petri dish containing fresh, chilled Hanks solution, and the hippocampi were separated from each other under binocular vision.

#### Vibratome (VT1200, Leica Mikrosysteme Vertrieb GmbH, Wetzlar) (second approach)

MEM solution was used during this preparation. The animal (P5) was first decapitated with scissors and the head was transferred to a Petri dish containing frozen MEM solution. The brain was carefully exposed on ice with small scissors and tweezers and directly attached to a cooled plate using tissue glue which was placed on the vibratome in cooled MEM solution. 350 µm thick brain sections were made along the sagittal plane. These brain sections were soaked up with a disposable pipette and transferred to a separate Petri dish containing cooled MEM solution. Hippocampi were then carefully dissected out under a microscope by removing the surrounding tissue except of a small part of the entorhinal cortex.

### 2.4 Cultivation

All steps necessary for cultivation were performed under sterile bench. A distinction is made between the cultivation of hippocampal slices in cell culture tubes and on inserts.

#### Roller tube

Prior to the acute preparation, 12 × 24 mm coverslips were placed in glass petri dishes and sterilized. One drop of collagen was then placed on each coverslip and spread over the total surface of the coverslip using a thin rubber tube. These were placed under UV light for 40 min to dry and stored sealed in the refrigerator overnight. Only vibratome sections were used for culturing coverslips in tubes. After preparation and just before placing the hippocampi on the coverslips, the coverslips had to be coated with plasma and thrombin. To do this, 10 µl of plasma was first added to each coverslip followed by 10 µl of thrombin. Both substances were then mixed on a small area and a hippocampus was placed on each. The coverslips were then set aside for 10 min to dry. Once the mixture is dried, an adherent surface forms so that the hippocampus does not detach (Humpel 2015). While the plasma-thrombin mixture was drying, the cell culture tubes were filled with 1 ml of BME medium heated in a heat bath. Finally, using tweezers, the coverslips with the hippocampal sections facing up were slid into the tubes and placed in the roll incubator, which rotated at a speed of 6 rpm. Cultivation was performed in an incubator at 36.5 °C.

#### Inserts (first approach)

The required inserts were transferred to 6-well plates filled with 1 ml MEM medium approximately two hours before the start of preparation and placed in the incubator or CO_2_ incubator. After all hippocampi were dissected out, further steps were performed with the aid of two spatulas. First, the supernatant fluid around the hippocampi was removed before four hippocampal slices per insert were placed on the membrane, one in each quadrant and with sufficient distance to avoid direct contact. The 6-well plates including the inserts were sealed with parafilm to prevent evaporation of the medium and placed in the incubator at 36.5 °C. The 6-well plates, which were stored in the CO2 incubator at 37 °C and 5% CO2 saturation, were not covered with Parafilm (Vlachos et al. 2012).

#### Inserts (second approach)

After dissecting the hippocampi one insert at a time was dipped into the MEM solution from below to moisten the membrane from below and then placed back in the package. Then 1 ml of MEM solution was pipetted from the side under the membrane and the hippocampi was placed on the membrane as previously mentioned. The insert was then placed into the 6-well plate, which was filled with 1 ml of MEM medium at the beginning of the experimental day. The 6-well plates, which were stored in the CO2 incubator at 37 °C and 5% CO2 saturation, were not covered with Parafilm (Vlachos et al. 2012).

On the day of preparation (day 0), the hippocampi were transferred to the coverslips of the rolling cultures or the inserts. The cultures were left to rest for the following day. The medium was changed every 2 days. Inserts of the second approach were left to rest for 18 days.

### 2.5 Preparing sections for histology

The cultured hippocampi had to be cut to a thickness of 30 µm. For this purpose, the microtome was cooled down to −25 °C. To create a base for cutting the cultures a spare piece of tissue (brain or liver) was first attached to the cutting plate using 30 % sucrose and frozen. Then, a straight surface was created with the cutting knife. One hippocampal culture at a time was transferred to a small coverslip with the aid of a brush and spread on the cooled base tissue. Then a brush was used to spread some 30% sucrose solution around the culture. Once everything was frozen through, 30 µm thick sections could be made.

### 2.6 Nissl staining

Hippocampal sections were mounted on chromalaungelatin-coated slides the day before Nissl staining and allowed to dry overnight at room temperature. Degreasing and dehydration of the tissue was performed by an ascending and descending alcohol series. First, an ascending alcohol series was performed in which the slides were briefly rinsed with distilled water and then immersed for 3 min each in 70% ethanol, twice 100% ethanol, isopropanol, and finally twice in xylene. This was followed by a descending alcohol series with xylene, isopropanol, twice 100% ethanol and finally 70% ethanol. The slides were then briefly rinsed again with distilled water and placed in the cresyl violet solution for approximately 4 min. After staining with cresyl violet, the slides were rinsed again with distilled water and then the ascending alcohol series was performed again as previously mentioned. After the excess xylene had been removed, the hippocampi could be mounted using the mounting medium DPX Mountant.

### 2.7 Immunohistochemical staining (IHC)

In the first step, hippocampal cultures used for IHC were washed three times in PBS for 10 min each. With the exception of PBS, all solutions were pretexted before use to ensure that all components were evenly distributed. Hippocampi were then transferred to a blocking solution of 10% normal Horse Serum in 0.2% PBS-Tx for 90 min at room temperature. The IHC was conducted as a 3-fold labelling, i.e., three primary antibodies were incubated simultaneously. The dilution medium consisted of 1% normal horse serum in 0.2% PBS-Tx and contained the following primary antibodies: mouse monoclonal antibody against NeuN diluted of 1:100, guinea pig polyclonal anti-GFAP diluted 1:500, and rabbit polyclonal anti-IBA1a diluted 1:250. Hippocampi were incubated overnight at room temperature with the primary antibody under gentle movement on the shaker & mixer. Next day, the hippocampi were again washed three times in PBS for 10 min each followed by incubation with the secondary antibodies, diluted in medium consisting of 1% normal Horse serum in 0.2% PBS-Tx: goat anti-mouse CY3 (1:500), donkey anti-rabbit Alexa 647 (1:500), and donkey anti-guinea pig Alexa 488 (1:250) for 90 min in the dark. All further steps were conducted under as little light exposure as possible. After incubation with the secondary antibody, the sections were washed again three times in PBS for 10 min each and then mounted on gelatinized slides. The slides were kept in the refrigerator overnight for air drying. The next day, the slides were rinsed with distilled water and allowed to dry again. Finally, the slides were covered with Dianova mounting medium free of air bubbles.

## 3 Results

In the following results section, different approaches for culturing postnatal hippocampal cultures are compared. A distinction is made between 5 methodological approaches:

1. Hippocampi were prepared in the Hanks solution, cut by vibratome without vibro-check function (first approach), the medium used was BME medium and the cultures were kept in an incubator without CO_2_ supply. Cultivation lasted 8 days.
2. Hippocampi were prepared in the Hanks solution, cut by vibratome without vibro-check function (first approach), the medium used was MEM medium and the cultures were kept in an CO_2_ incubator. Cultivation lasted 8 days.
3. Hippocampi were prepared in the Hanks solution, cut by a chopper, the medium used was MEM medium and the cultures were kept in an CO_2_ incubator. Cultivation lasted 8 days.
4. Hippocampi were prepared in the Hanks solution, cut by vibratome with vibro-check function (preparation was done like the first approach), the medium used was MEM medium and the cultures were kept in an CO_2_ incubator. Cultivation lasted 2 weeks.
5. Hippocampi were prepared in the MEM solution, cut by vibratome with vibro-check function (second approach), the medium used was MEM medium and the cultures were kept in an CO_2_ incubator. In addition, a part of the entorhinal cortex stayed attached to the hippocampus and the cultivation lasted 18 days.

### 3.1 Histological comparison of directly fixed hippocampi

In order to get an impression of the condition of the hippocampi, some were cut during preparation and directly fixed to determine if there was any damage to the tissue due to the mechanical influence of cutting. The directly fixed (df) hippocampal slices were not cultured and show the condition of the tissue before cultivation. Df hippocampal sections were fixed directly in 4% PFA after dissection and then stored in 30% sucrose solution until further processing. In Figure 1 the upper image shows the staining of hippocampi sectioned through the vibratome without vibro-check, the middle image shows hippocampi sectioned through the chopper and the lower image shows hippocampi sectioned through the vibratome with vibro-check. All images show undamaged structure of the GD and CA regions. Also, the CA3 region can be clearly seen by the more loosely packed pyramidal cells, compared to the densely packed CA1 region. The vibratome without vibro-check function was used in the first and second cultivation methods. It was then replaced by the chopper to see if it had a positive effect on the survival of the neurons, as the preparation with the chopper was faster in time. Finally, the vibratome with vibro-check function was used to exert the least mechanical influence on the cells (Zimmermann et al. 2009).

**Figure 1.**
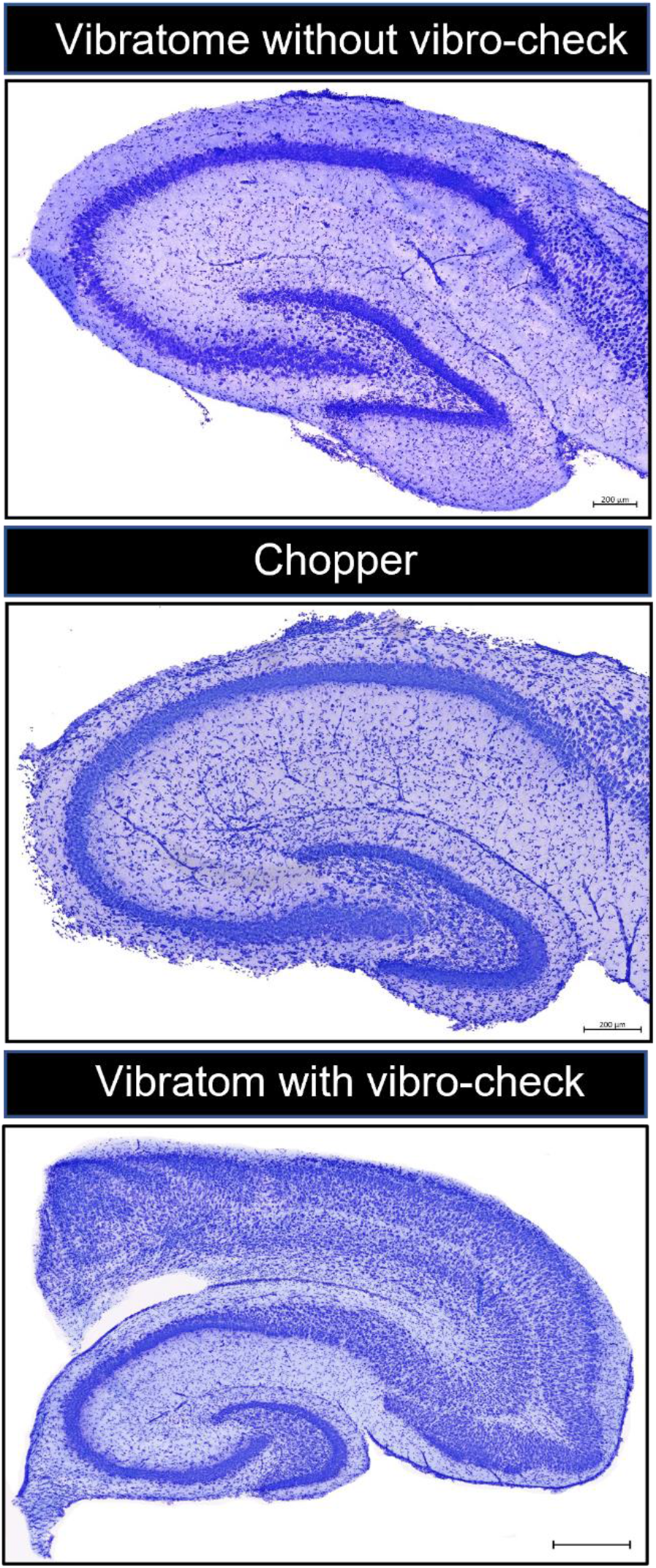
Comparison of directly fixed (df) hippocampi sliced through the vibratome without vibro-check function (top), through the chopper (middle) and through the vibratome with vibro-check function (bottom). All hippocampi show complete structure of dentate gyrus and cornu ammonis. Hippocampal cultures cut with the vibratome with vibro-check stayed attached to a part of the entorhinal cortex. df = directly fixed; GD = dentate gyrus; CA = cornu ammonis. Scale = 200 µm (top and middle image) and 500 µm (bottom image).

### 3.2 Histological analysis of the roller tube cultures

Initially, cultivation by roller tubes was tested. It was documented in time how the tissue changed. Figure 2 shows the result of Nissl staining of the roll tube cultures. The top image shows a hippocampal section after five days in culture using roller tubes. It can be seen that the hippocampus has lost its original shape. The structure appears torn and incomplete. The only clearly recognizable structure is that of the GD. CA3 and CA2 are no longer present and CA1 cannot be visually distinguished from the rest of the cell cluster. The middle and bottom images show the result of culturing after 10 and 15 days, respectively. In both cultures, no shape or structure of the hippocampus can be seen anymore. Day 15 also shows a washing away of the cells from the tissue association

**Figure 2.**
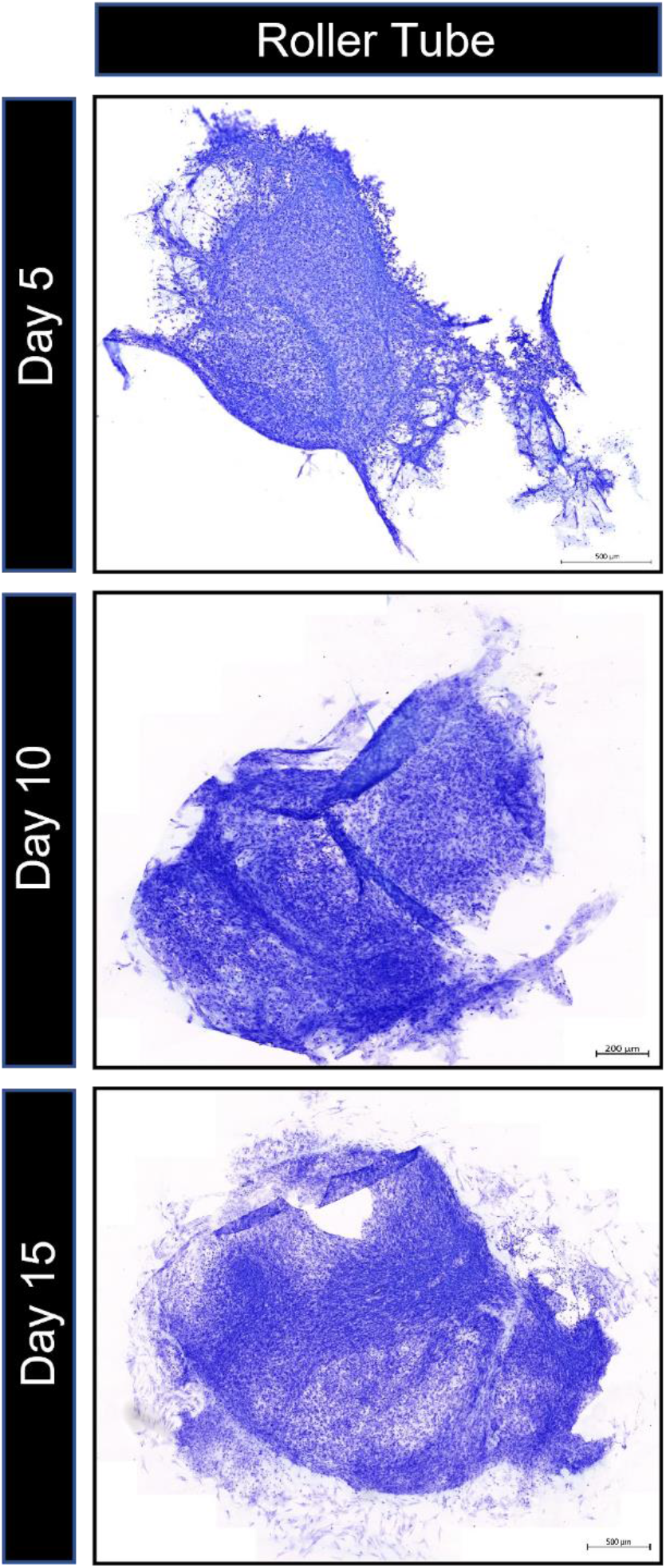
This figure compares the hippocampal slices after 5, 10 and 15 days in culture using roller tubes. After five days, the hippocampus no longer has its original shape. The only structure that can be seen is the dentate gyrus. After 10 and 15 days, no hippocampal shape can be seen, and no structures can be made out anymore. The image at Day 15 shows a washing away of cells from the tissue association. Scale = 500 µm.

### 3.3 Immunohistochemical comparison of different cultivation methods

Figure 3 compare the effects of the first three different cultivation methods mentions before on neurons, astrocytes and microglia. Immunhistochemically, neurons were stained by NeuN (red), astrocytes by GFAP (green), and microglia by IBA1a (blue). The GFAP staining in the 2^nd^ method cannot be shown due to overexposure of the channel because of a defect of the antibody. In the 1st method, overexpression of GFAP can be seen and slight expression of NeuN and IBA1a. In the 2nd method (GFAP cannot be shown), few NeuN-expressing neurons are seen. The 3rd method shows lighter expression of GFAP and stronger expression of IBA1a. Neurons are seen in the dentate gyrus (GD) and cornu ammonis region 1 (CA1).

**Figure 3.**
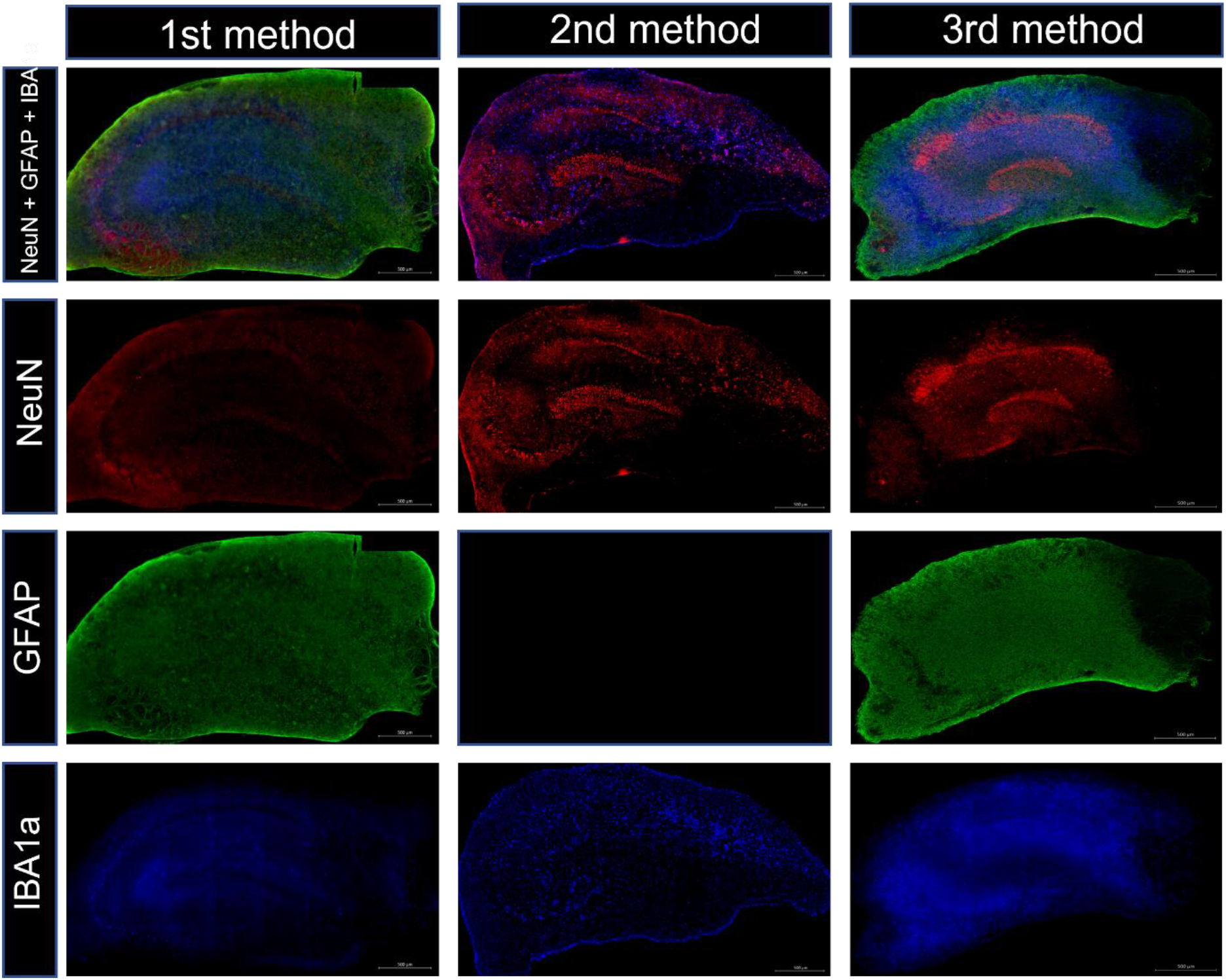
This figure shows the immunohistochemical results of the hippocampal slices cultured after the 1st, 2nd and 3rd method. Neurons were stained by NeuN (red), astrocytes by GFAP (green) and microglia by IBA1a (blue). In the 1st method, overexpression of GFAP in can be seen and slight expression of NeuN and IBA1a. In the 2nd method (GFAP cannot be shown due to severe overexposure), few NeuN-expressing neurons are seen. The 3rd method shows lighter expression of GFAP and stronger expression of IBA1a. Neurons are seen in the dentate gyrus (GD) and cornu ammonis region 1 (CA1). GD = dentate gyrus; CA = cornu ammonis. Scale = 500 µm.

### 3.4 Histological comparison of cultured hippocampi with and without an intact cortex dressing

Figure 4 shows an exemplary example of a culture after two weeks of cultivation without an intact cortex association. The hippocampus was sectioned by the chopper. The basic shape is unchanged. The tissue looks torn and shows cell loss in several areas. The only faintly discernible structure is that of the dentate gyrus. The region of the CA can no longer be identified due to the lack of tissue. In general, the hippocampus appears densely packed and overgrown with cells. Figure 5 shows the result after two weeks of cultivation with the cortex intact. The preparation was performed using the vibratome with vibro-check function. The top row shows overview of the whole structure. The cortex appears slightly torn and pitted. The hippocampi themselves show no cell loss. Looking at the hippocampal sections of the bottom row, both the GD and CA regions can be seen clearly. In general, the hippocampi appear densely packed and overgrown with cells, making the hippocampal formation stand out only faintly.

**Figure 4.**
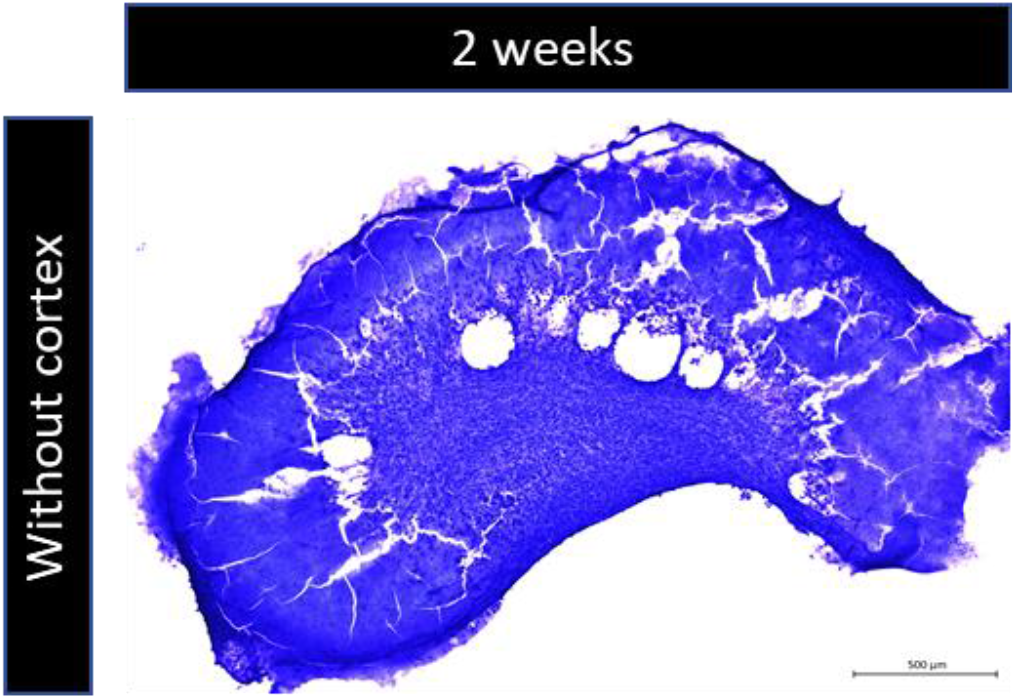
The image shows Nissl staining of a hippocampus without an intact cortex association after two weeks of cultivation. The basic structure of the hippocampus is unchanged. The tissue itself shows tears and holes. The only discernible region is that of the dentate gyrus. The cornu ammonis region can no longer be identified. GD = dentate gyrus. Scale = 500 µm.

**Figure 5.**
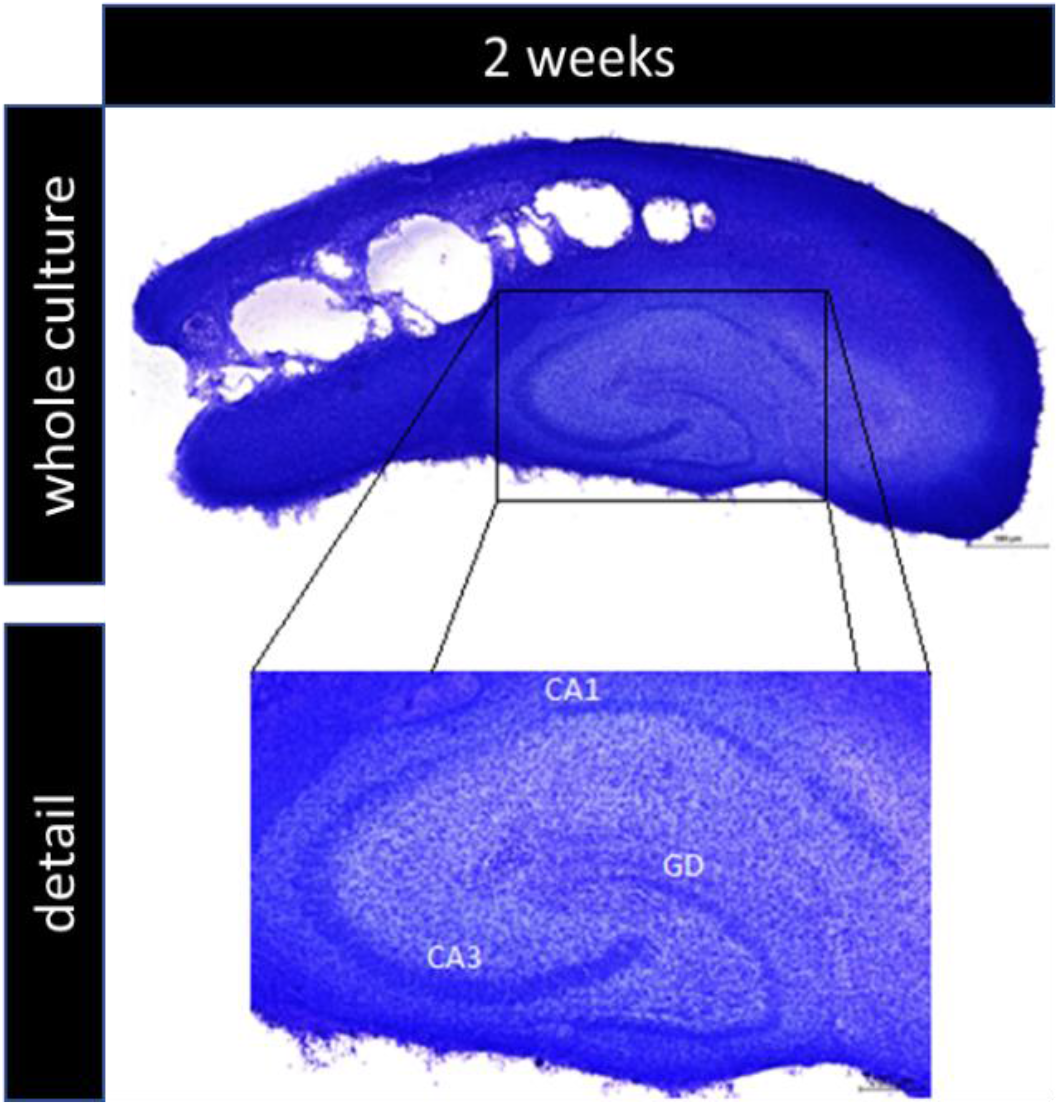
The top row shows the staining of the entire culture after 2 weeks. The cortex shows holes and cracks, whereas no cell loss is seen in the hippocampal formation. The bottom row highlights the section of the hippocampus. The dentate gyrus and cornu ammonis structures are complete. GD = dentate gyrus; CA = cornu ammonis. The scale is 500 µm for overview of the whole culture and the sections of the hippocampi are 200 µm.

### 3.5 Immunohistochemical analysis of cultured hippocampi with attached cortex after two weeks (4^th^ method)

The following Figure 6 shows the image of the immunohistochemical staining of the hippocampi with attached cortex after two weeks of cultivation. The vibratome with vibro-check function was used for the preparation. Neurons were stained by NeuN (red), astrocytes by GFAP (green), and microglia by IBA1a (blue). The figure shows the all three channels superimposed. NeuN staining shows loss of neurons, both in the GD and CA regions. Only in the GD isolated neurons can be detected. When looking at GFAP and IBA1a staining, high expression can be seen throughout the hippocampus. Immunohistochemical comparison to hippocampal sections after two weeks of cultivation without cortex cannot be shown because the tissue did not withstand the mechanical influence of immunohistochemical staining. The condition of the tissue can be seen from Figure 4. When the cultures were viewed with the naked eye, it could be seen that the original hippocampal section had thinned out. The cultures without cortex appeared translucent compared to the cultures with cortex.

**Figure 6.**
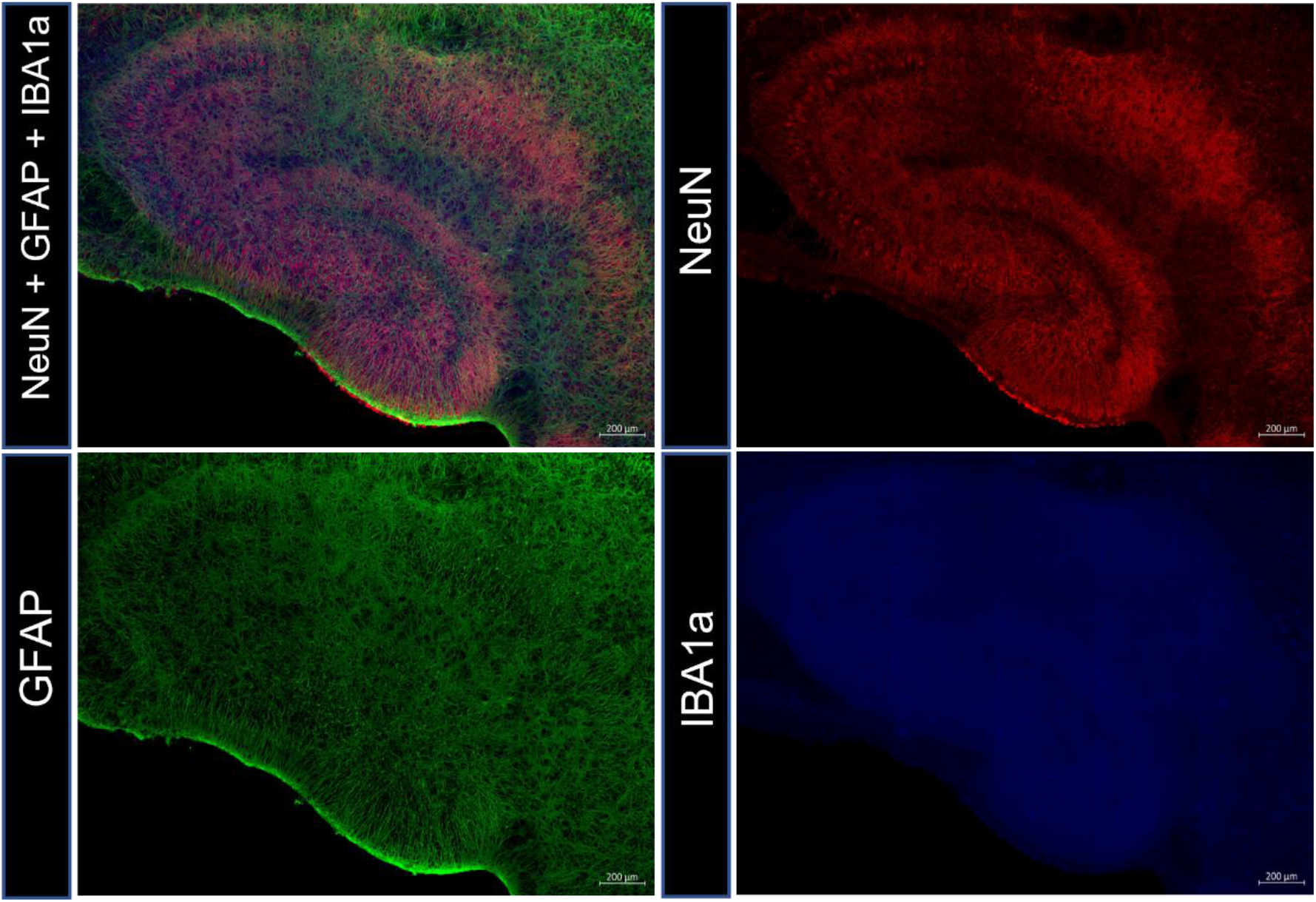
Shown is the immunohistochemical image of the hippocampi with intact cortex after two weeks of cultivation. Neurons were stained by NeuN (red), astrocytes by GFAP (green) and microglia by IBA1a (blue). The image shows all three channels superimposed at the top left and all single channels. Only a few isolated neurons be detected. In addition, a high expression of GFAP and IBA1a positive cells can be seen. Scale = 200 µm.

### 3.6 Immunohistochemical analysis of cultured hippocampi with attached cortex (5^th^ method)

The cultured hippocampi were allowed to rest in CO_2_ incubator for 18 days after dissection. The preparation was performed according to the second approach (see 2.4 Inserts (second approach)). Figure 7 shows the immunohistochemical staining of the results after the fifth method. Neurons were stained by NeuN (red), astrocytes by GFAP (green), and microglia by IBA1a (blue). The image does not show overexpression of IBA1a and GFAP as in the previous methods. Intact neurons are seen in all four regions of the hippocampus (GD, CA1, CA2, and CA3). Furthermore, the cultured tissue withstood the mechanical influence of the microtome, and more sections could be obtained from one hippocampus than with the previously performed methods.

**Figure 7.**
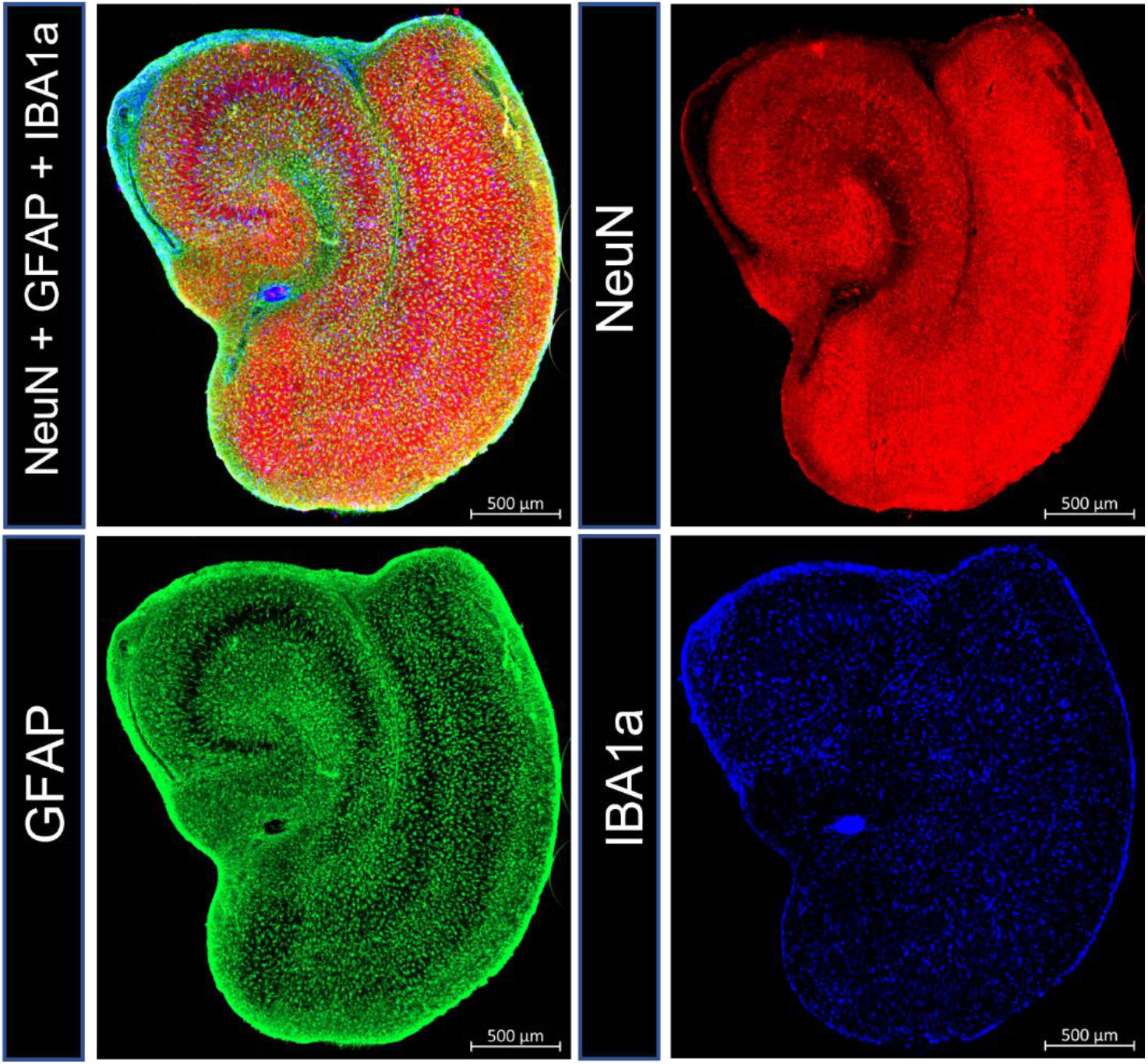
Figure 7 shows the result of immunohistochemical staining according to the preparation method (inserts-second approach) and the fifth cultivation method. The staining was performed after a resting period of 18 days in the CO_2_ incubator. Neurons were stained by NeuN (red), astrocytes by GFAP (green) and microglia by IBA1a (blue). The first image shows all 3 channels superimposed. Neurons are well visible throughout the hippocampal formation and no overexpression of GFAP and IBA1a positive cells is seen. This can also be seen when looking at the individual channels. Especially in the individual channel GFAP, it can be clearly seen that the GD and CA region is not overgrown with glial cells. GD = dentate gyrus; CA = cornu ammonis. Scale = 500 µm.

## 4 Discussion

### 4.1 Loss of hippocampal structure in roll cultures

The roll culture method was first described by Hogue in 1947 (HOGUE, 1947) and the cultivation of brain slices in cell culture tubes was modified by Gähwiler in 1981 (Gähwiler 1981a, 1981b; Humpel 2015). By flattening the tissue in this method, the neurons arranged themselves in a monolayer, which made it possible to view them using phase-contrast microscopy. However, it was also observed that detachment or dissolution of the cultures occurred (Gähwiler 1981a, 1981b; Humpel 2015). In this work, the hippocampi were initially cultured in cell culture tubes, however, the previously mentioned effects occurred. Figure 2 shows structural alteration of the hippocampi. It can be clearly seen that degeneration of the tissue occurred. The hippocampal structure is no longer present in all three examples. The cells appear scattered and no longer belong to the tissue association. Furthermore, there was a loss of cultures during cultivation, on the one hand the cultures detached from the coverslips and on the other hand the cultures also dissolved (Humpel 2015). One reason may be the mechanical influence, as the cell culture tubes were in a rolling incubator that rotated at 6 rpm throughout. Another reason may be the lack of CO2 supply. The pH of the medium is constantly regulated by the CO2 supply, so a less than optimal atmosphere can cause the pH to become more acidic or basic (Autrup et al. 1978; Dame et al. 2010). The higher the CO2 content, the lower the pH of the medium (Schmitz 2011). This can lead to an impairment of cell activity or even to cell death (Gourine et al. 2010; Ozawa et al. 2006).

### 4.2 Acceleration of preparation has positive effects on neuron survival

In 1991, the rolling culture method was modified by Stoppini et al. (Stoppini et al. 1991). They used semipermeable membranes, which prevented the cultures from detaching and being lost. Also, the mechanical influence on the cultures was reduced, as the cultures were no longer in constant motion, but only moved at the time of a medium change (Humpel 2015; Stoppini et al. 1991). The medium was pipetted under the so-called inserts so that the membrane was in contact with the medium from below, but the cultures above were not surrounded by medium (Humpel 2015; Humpel 2018; Stoppini et al. 1991). To eliminate the problems observed with rolling cultures, this work switched to semipermeable membranes. This resulted in a significant improvement in tissue structure, which was maintained even after several days of cultivation. Furthermore, detachment and dissolution of the cultures, as described by Stoppini et al. and Humpel (Humpel 2015; Stoppini et al. 1991), also did not occur. However, the problem of gliosis occurred, which Stoppini already mentioned in 1991 in connection with his cultivation on a membrane. Microglia respond rapidly to tissue injury, in epileptic seizures, or inflammatory processes in the brain (Feng et al. 2019), whereas astrocytes respond with hyperplasia, hypertrophy, and increased GFAP immunoreactivity (Tarasov et al. 2019).In this work, there was increased growth of astrocytes and microglia and loss of neurons, which may be explained by preparation-related tissue injury. To prevent the occurrence of gliosis, the methodological approach was optimized several times, also with the aim of improving the general condition of the tissue and reducing the loss of neurons.

Figure 3 presents the results of the different methods. The first method, using the vibratome without vibro-check function, BME medium, and incubator without CO2 supply, shows increased expression of GFAP and IBA1a and decreased expression of NeuN. Several reasons may account for this. On the one hand, it could be due to the vibratome without vibro-check function. The vibro-check function serves to compensate for the vibrations of the razor blade generated by the device and thus protect the tissue from mechanical damage (Zimmermann et al. 2009). If this is not the case, the mechanical impact can lead to cell death. Also, cutting took a long time, approximately 30 minutes per brain, so there was a risk that the tissue was not sufficiently and consistently cooled before culturing. Another reason for low NeuN expression could be the lack of CO2 supply, which affects the pH of the medium and can affect cell activity and lead to cell death (Dame et al. 2010; Gourine et al. 2010). On the other hand, improper composition and pH adjustment of the medium can also have effects on culture survival (Chen et al. 2006; Higashino et al. 2018; McAdams et al. 1997; Teo et al. 2014).

In the second method, the vibratome without vibro-check function was also used, but the BME medium was replaced by the MEM medium and the incubator without CO2 supply was replaced by the CO2 incubator. Cultivation with the MEM medium in the CO2 incubator showed less development of gliosis. The MEM medium is similar to the BME medium previously used in the first method but contains few substantially different ingredients. MEM is a modification of BME and contains a higher concentration of amino acids (Schmitz 2011). Amino acids are essential for the organism and accordingly must be considered during cultivation to ensure cell growth. L-glutamine is the only amino acid that can cross the blood-brain barrier (Kawai et al. 1999) and is necessary for cell metabolism. However, L-glutamine does not remain stable for long at a temperature of 37 °C, so glutamax acts as a substitute in the MEM medium. It is more stable than L-glutamine and has a growth-promoting effect on cells. In addition, sodium bicarbonate is used in the MEM medium, which supplies the cells. Its buffering capacity prevents the metabolic products of the cultures produced in the medium from over-acidifying it. Furthermore, the contained HEPES buffer supplements the buffer capacity of the medium (Schmitz 2011). The ingredients of the MEM medium protect the cultures more strongly from pH fluctuations and metabolic waste products, so that the tissue is optimally supplied, which can be seen in the lower growth of the glial cells. However, low NeuN expression is still seen. This suggests that too much mechanical influence was exerted on the tissue by the vibratome without vibro-check function, resulting in degeneration of the cells, since the vibratome was the only component that was not replaced.

Another methodological adjustment was the preparation with the chopper in the third method. Previously, the vibratome was used without the vibro-check function, which means that the vibrations of the blade may have caused the tissue to receive additional mechanical impact (Zimmermann et al. 2009), resulting in neuronal death. Cutting the hippocampi with the chopper was faster in time (approximately 10 minutes per brain) than with the vibratome without the vibro-check function, so there was no risk of breaking the cold chain. Also, the sections could be in culture more quickly, allowing the tissue to recover after dissection. The results show a higher number of neurons, which means that the faster preparation had a positive effect on neuron survival.

### 4.3 Longer cultivation stabilizes the tissue of cultures

To get a better overview of the progress of the cultivation a time series was recorded over an eight-day period (Data not shown). Cultures were fixed daily to observe changes of neurons, astrocytes and microglia in more detail and to see how the tissue structure changes over time. It was clear that the structure of the hippocampus stabilized over time, as seen by the reduction of tears and holes in the tissue, however the growth of astrocytes and microglia also increased. According to Humpel 2015, cultures should remain in culture for at least two weeks to allow the tissue to recover from dissection, the cultures to attach to the membrane, and the sections to have time to thin out (Humpel 2015; Humpel 2018). Therefore, it can be explained that the hippocampi showed a better general optical condition after at least eight days of cultivation than only a few days after dissection. Axotomy may be another reason for stronger staining of glial cells and weaker staining of neurons (Humpel 2015; Richardson et al. 2009). Dissecting out the hippocampi severed all afferent and efferent nerve fibers. This can lead to Waller degeneration, meaning that the distal portion of the axon cannot continue to be supplied (Conforti et al. 2014; Llobet Rosell und Neukomm 2019). Looking at Figure 3, it is noticeable that there was lower expression of NeuN in all regions of the hippocampus. This suggests that removal of the entorhinal cortex and thus damage to the tractus perforans resulted in Waller degeneration and that this had an impact on neuron survival.

### 4.4 Cultivation with intact cortex dressing prevents tissue degeneration

Further cultivation of hippocampal slices was extended to two weeks. Initially, chopper slices were cultured for two weeks without entorhinal cortex. The result is shown in Figure 4. However, by the end of the two-week cultivation, the tissue showed cracks and was partially dissolved. The tissue had been too fragile for immunohistochemical staining. This reinforces the suggestion that the consequence of axotomy is Waller degeneration (Conforti et al. 2014; Llobet Rosell und Neukomm 2019). Embryonic and postnatal brains in particular are sensitive and rely on supply or projections from the entorhinal cortex during development (Humpel 2018). Because dissection occurred at P10, removal of the entorhinal cortex resulted in long-term damage to cultures because postnatal sections no longer received projections. This explains why no damage or loss of neurons was seen in the df hippocampal cultures but later in the culturing process. Subsequently, cultures were made in which the hippocampus was connected to the entorhinal cortex during culture. In addition, hippocampal slices were cut using the vibratome with vibro-check function. The vibro-check function exerts less mechanical influence on the tissue, which in turn protects the tissue from injury (Zimmermann et al. 2009). Figure 5 shows the result of Nissl staining after two weeks of cultivation. The cortex appears torn and partially dissolved. In comparison, however, no damage is seen in the hippocampus. The hippocampal formation is preserved and visible, however the tissue appears overgrown compared to df sections. Compared to the hippocampi cultured for two weeks without intact cortex dressing, the hippocampi with intact cortex dressing withstood immunohistochemical staining. Looking at the immunohistochemical staining in Figure 6, it is noticeable that despite the intact tissue, high GFAP and IBA1a expression is seen and NeuN expression is weaker. The results suggest that preserving the connections to the entorhinal cortex and thus avoiding axotomy has positive effects on the state of the tissue by slowing down degeneration of the cells (Conforti et al. 2014; Llobet Rosell und Neukomm 2019). However, at this stage, there is still the problem that the hippocampi present a loss of neurons compared to the perfused and df sections, and there is an increased growth of astrocytes and microglia.

### 4.5 Optimal combination of methodical procedures

In the last cultivation method (Fig. 7), the combination of several factors had a positive effect on neuronal preservation and glial growth in hippocampal cultures. Firstly, the Hanks solution was replaced by the MEM solution during the preparation. In contrast to the Hanks solution, the MEM solution contained all components that are contained in the MEM medium, which meant that the sections were not exposed to two different solutions (see 4.2 for the effects of the ingredients). In addition, the pH of the MEM solution was previously adjusted, and the solution was not gassed with carbogen, which could result in no pH fluctuations, which may have a negative effect on cell survival in the hippocampus (Schmitz 2011; Gourine et al. 2010; Ozawa et al. 2006). Furthermore, the preparation was faster because the brain was not immersed in cold preparation solution beforehand and then cut to size. Immediately after exposure, the rat brain was attached to a cooled plate with tissue glue, with the cortex facing down. This was placed directly into the cooled chamber of the vibratome filled with MEM solution and cut. This has the advantage that there is no interruption of the cooling chain. Another point that was handled differently is that only the ventral hippocampus was used for culturing, as there are functional differences between the ventral and dorsal hippocampus (Fanselow und Dong 2010). This allows change to be compared more accurately. To prevent Waller degeneration, part of the entorhinal cortex remained connected to the hippocampus (Conforti et al. 2014; Llobet Rosell und Neukomm 2019). This results in a shorter preparation time, which allowed the sections to be transferred to the membranes more quickly and thus placed in the cultivation medium in the CO2 incubator more quickly. By resting for 18 days with 3 media changes per week, the tissue had time to regenerate from the preparation, which stopped the loss of neurons and prevented the overgrowth of astrocytes and microglia (Humpel 2015). When handling the tissue after culturing, the positive effect on the tissue was confirmed. The sections were more robust and more 30 µm thick sections could be obtained when cutting with the microtome.

### 4.6 Outlook

In this research, different methods were compared to find the optimal organ cultivation method for postnatal hippocampal cultures. The change from rolling cultures to semipermeable membranes showed a substantial improvement in terms of the general condition of the tissue and the preservation of the hippocampal structure (Humpel 2015). Further downstream, faster preparation was shown to have positive effects on neuron survival and reduced growth of astrocytes and microglia (Humpel 2018). However, cultivation in a CO2 incubator also improved the general condition of the cultures (Gourine et al., 2010). Through the recorded time series, it was shown that longer cultivation stabilized the tissue (Data not shown) (Humpel 2015; Humpel 2018). The subsequent two-week cultivation provided good insight into whether it is useful and necessary to culture postnatal hippocampal cultures with the cortex intact. The tissue of the hippocampi with intact cortex dressing was much more stable in contrast to the hippocampi without intact cortex dressing and did not show cracks and dissolved tissue (Vlachos et al. 2012). By considering all the factors that had a positive effect and additionally adjusting the preparation solution, an optimal result was obtained. In summary, it can be concluded that various aspects have an influence on successful cultivation. Not only the cultivation method, but also the type of preparation, the desired structure, the medium and its pH, and a CO_2_-containing environment play a role in a promising organ culture.

